# Hybrid Discrete-Continuum Cellular Automaton (HCA) model of Prostate to Bone Metastasis

**DOI:** 10.1101/043620

**Authors:** Arturo Araujo, David Basanta

## Abstract

Prostate to bone metastases induce a “vicious cycle” by promoting excessive osteoclast and osteoblast mediated bone degradation and formation that in turn yields factors that drive cancer growth. Recent advances defining the molecular mechanisms that control the vicious cycle have revealed new therapeutic targeting opportunities. However, given the complex temporal and simultaneous cellular interactions occurring in the bone microenvironment, assessing the impact of putative therapies is challenging. To this end, we have integrated biological and computational approaches to generate an accurate model of normal bone matrix homeostasis and the prostate cancer-bone microenvironment. The model faithfully reproduces the basic multicellular unit (BMU) bone coupling process and introduction of a single prostate cancer cell yields a vicious cycle that is similar in cellular composition and pathophysiology to models of prostate to bone metastasis.

## I. INTRODUCTION TO THE TYPE OF PROBLEM IN CANCER

Prostate cancer metastases in the bone induces osteolytic and osteoblastic lesions that are intensely painful and greatly contribute to the morbidity associated with the disease [1]. Our current understanding bone metastatic prostate cancer growth is encapsulated by the “vicious cycle” paradigm in which prostate cancer cells manipulate bone forming osteoblasts and bone resorbing osteoclasts to yield growth factors and space for expansion and growth [2]. Currently bone metastatic prostate cancer is incurable and only by understanding the dynamics of molecular and cellular interactions driving the cancer can we hope to develop curative strategies. We posit that integrating relevant biological parameters and models with a computational modeling approach can be a powerful means with which to address this clinically significant problem. To this end, we developed a novel computational model of the prostate cancer bone microenvironment [3]. Our research suggests that a key advantage of this kind of biologically relevant computational model is that it opens a window into the inner dynamics of the cells, their heterogeneity, their interactions and their behaviors across time and space; features that are difficult to determine using traditional experimental approaches.

In our computational model, each cell is represented individually using an agent-based Hybrid Discrete Cellular Automata (HCA) approach [4,5] and realistic biological behavior emerges from the interactions of the cells (Figure 1). The system recapitulates the normal bone modeling process and the delicate balance between bone regenerating cells; mesenchymal stem cells (MSCs), precursor osteoblasts (pOBs), adult osteoblasts (aOBs) and bone resorbing cells; precursor osteoclasts (pOCs) and osteoclasts (OCs). Critical to the balance is the bioavailability of factors such as transforming growth factor β (TGFβ), receptor activator of nuclear kappa B ligand (RANKL) and bone-derived factors generated by osteoclasts. The introduction of a metastatic prostate cancer cell into the system that expresses the TGFβ receptor and the ligand (albeit at lower concentrations that that derived from the bone) results in the recruitment of MSCs, an unregulated pOB expansion and the influx of osteoclast precursors. This in turn leads to increased osteoclast formation and bone destruction. The release of bone-derived nutrients and sequestered growth factors from the bone matrix, such as TGFβ, promotes the survival and growth of the metastatic prostate cancer cells, thus perpetuating the vicious cycle that mimics the pathophysiology of the human clinical scenario (Figure 1).

**Figure 1.**
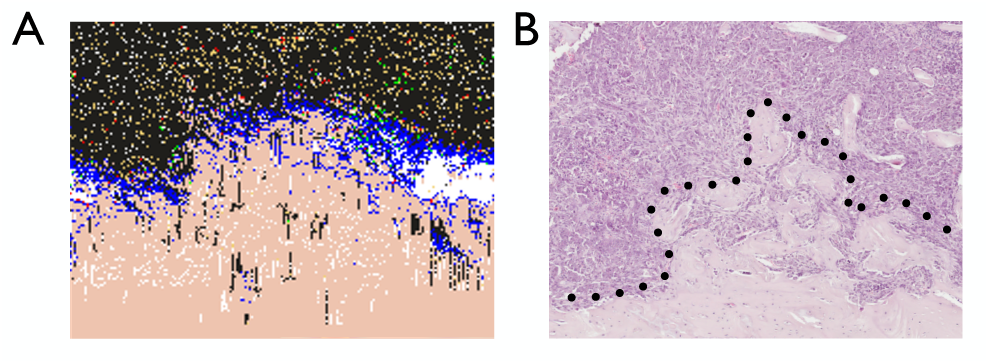
Computational model output of the prostate cancer-bone microenvironment is consistent with histological analysis of the in vivo microenvironment. A-B, Analysis of the of the computational output (A) of the prostate cancer-bone microenvironment is similar to that of and in vivo prostate cancer to bone metastasis model (B).

Some of the key novel insights derived from this computationally-led research are 1) the phasic nature of the vicious cycle 2) that MSCs significantly contribute to prostate cancer induced osteogenesis and, 3) TGFβ is a critical mediator of communication between each of the cellular compartments. Comparison of the computational model outputs with an in vivo model of the disease confirms the usefulness of the computational model (Figure 2).

**Figure 2.**
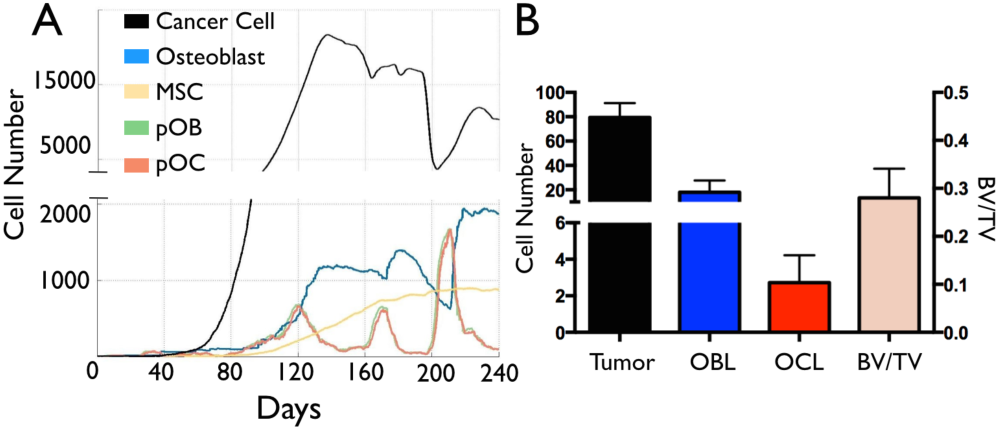
Analysis of the changes in cell population over time in the computational simulation of the prostate cancer bone microenvironment. A. Dynamic changes in the environmental cell populations can be observed and studied. B. These outputs inform ongoing experiments at the Moffitt Cancer Center’s Lynch Lab on the inner dynamics of the system.

In addition to genetic mutations, it has been established that the surrounding tumor-microenvironment is also a major driver of cancer evolution [6]. A major feature of our computational model is the response of the bone microenvironment to the cancer cells. Therefore, by allowing the model to include cancer heterogeneity and allowing the population of tumor cells to evolve we posit that it is a significant predictive tool that will ultimately aid oncologists in choosing the combination and sequence of therapies to administer to individual patients.

## II. LLUSTRATIVE RESULTS OF APPLICATION OF METHODS

Emerging results show how computational modeling can provide key insights into the dynamics of bone metastatic prostate cancer. Importantly, the model reveals distinct phases of osteolytic and osteogenic activity; a critical role for mesenchymal stromal cells (MSCs) in osteogenesis and temporal changes in cellular composition (Figure 2). A major motivation for building such models is the ability to use them to assess the efficacy of current and putative targeted therapies. To determine the robustness of the model, we applied currently utilized bisphosphonate and anti-RANKL therapies to established metastases. At 100% efficacy, bisphosphonates inhibited cancer progression while, in contrast to clinical observations; anti-RANKL therapy completely eradicated the metastases. Lowering the efficacy of the anti-RANKL therapy yielded clinically similar results suggesting that better targeting or dosing could improve patient survival. In this regard, we have successfully modeled the impact of therapies of the current standard of care for bone metastatic prostate cancer such as the anti-RANKL drug, denosumab [3],

## III. QUICK GUIDE TO THE METHODS

The computational model is based on the Hybrid Discrete-Continuum Cellular Automaton paradigm (HCA). In it, tumor cells live in a grid of 200x50 points representing 2 x 0.5 mm^2^ of the bone. A major advantage of the HCA is in its intimate interconnection with experimental data, where the model and the experiments inform each other. This increases the accuracy of the model abstractions and connectivity of the basic elements, which yields reliable and biologically relevant emergent behaviors. The core of the computational model recapitulates the normal BMU program. This involves, 1) retraction of the osteoblasts from the bone surface and generation of a canopy, 2) division of MSCs to generate RANKL expressing osteoblast precursors that, 3) recruit and promote the activation of bone resorbing osteoclasts and, after resorption and osteoclast apoptosis, 4) osteoblasts undergo differentiation and repair/restore the resorbed area (9, 10). To model this, we have focused on understanding the role and behavior of the key regulators of the BMU dynamics. The principal cellular players were explicitly modeled as agents in a grid following specific rule sets in a physical microenvironment.

The model considers 7 different cell types, 6 natural residents in the BMU microenvironment: Osteoblasts (Ob), Osteoclasts (Oc), precursor Ob, precursor Oc, Mesenchymal Stem Cells (MSCs), bone; as well as prostate tumor cells (PCa), parameterized with the help of biological experiments, capable of recruiting MSCs and producing their own TGFβ. The dynamics of the different cell types are mediated by the microenvironmental elements RANKL, TGFβ and bone derived factors (BDFs); characterized by partial differential equations that are subsequently discretized and applied to a grid. TGFβ is produced by bone destruction (α_β_B_i,j_) and cancer cells (α_c_C_i,j_) in proportion to the local TGFβ concentration, with natural decay of the ligand (σ_β_T_β_); ensuring the density never exceeds a saturation level, m_0_. TGFβ has pleiotropic effects on osteoblasts, osteoclasts and metastatic prostate cancer cells. TGFβ regulates osteoblast function with mutations in TGFβ signaling leading to severe bone phenotypes such as osteogenesis imperfecta (7). Low concentrations of TGFβ stimulate osteoclastogenesis but high concentrations inhibit the process; illustrating the biphasic effects of this growth factor even on the same cell type (8, 9). Our group and others have shown that TGFβ supports tumor survival and growth by activating TGFβ receptors on the tumor cell surface (10-12). TGFβ is governed by the following differential equation:

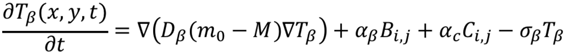

RANKL R_L_ is produced by precursor Osteoblasts, α_L_ O_i,j_, in proportion to the local RANKL concentration, with natural decay of the ligand, σ_L_ R_L_; ensuring the density never exceeds the saturation level n0. The concentration of RANKL is determined by:

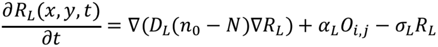

Factors F_B_ are released by bone destruction, α_B_B_i,j_, in proportion to the local Factor concentration, with natural decay of the factors, σ_B_F_B_; ensuring the density never exceeds the saturation level p_0_. As such, the dynamics of the bone related factors is calculated through:

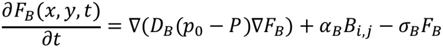

Periodic boundary conditions were considered only for the left and right sides of the microenvironment, while no-flux boundaries were imposed on the top and bottom of the two dimensional grid.

In the model, homeostasis is normally maintained in the absence of cancer cells and a vicious cycle emerges when cancer cells are introduced. Focusing on capturing the identities of the key players, behavior naturally emerges from the interactions of the microenvironmental components. In conclusion, we have generated an accurate computational model that can be tailored for the rapid assessment of putative therapies and the delivery of precision medicine to patients with prostate to bone metastases

## ACKNOWLEDGEMENT

We would like to thank Leah Cook, Conor Lynch and the IMO department at Moffitt Cancer Center for the scientific and moral support for this interdisciplinary research.

